# CRB2 controls primary ciliogenesis in association with PAR6α, HOOK2 and RAB8A

**DOI:** 10.1101/830166

**Authors:** Magali Barthelemy-Requin, Dominique Massey-Harroche, Stéphane Audebert, Jean-Paul Borg, Jan Wijnholds, André Le Bivic

## Abstract

Ciliogenesis is an essential process in animal development and physiology and when affected, the origin of severe pathologies called ciliopathies. CRB2 has been recently implicated when mutated in several pathologies such as retina degeneration and a ciliopathy-like syndrome. Here, we have investigated the role of CRB2 in primary ciliogenesis and showed that CRB2 depletion blocks the formation of the ciliary vesicle after activation of the mother centriole, indicating a defect in either transport or anchoring of the ciliary vesicle. CRB2 is accumulated in intracellular vesicles in the vicinity of the centrosome and we showed here that CRB2 forms a complex made of HOOK2 and PAR6α, two proteins important for primary ciliogenesis or centrosome integrity. In addition, CRB2 interacts with RAB8A which over-expression rescued CRB2 knockdown. All together, these findings indicate that CRB2 regulates a membrane transport step essential for the formation of the ciliary vesicle.

## INTRODUCTION

Defects in the formation or function of primary cilium result in severe pathologies such as ciliopathies and retina degeneration in human and many genes have been involved in this process. Among these genes, *CRB2 (Crumbs2)* has been recently implicated in several human pathologies reminiscent of ciliopathies. Indeed, *CRB2* mutations have been associated in the last years with a growing body of defects that have been regrouped recently in the *CRB2*-related syndrome of poor prognosis resembling congenital Finnish nephrosis (Jaron et al., 2016; Lamont et al., 2016; Slavotinek et al., 2015). This *CRB2*-related syndrome is characterized by cystic renal disease and cerebral ventriculomegaly (Slavotinek, 2016) suggesting that ciliogenesis or ciliary functions are compromised. To support this hypothesis, it has been shown that in the zebrafish, a Crb2b isoform is essential for the elongation of the inner segment of photoreceptor (PR) and of the primary cilium of kidney cells (Hazime and Malicki, 2017; Omori and Malicki, 2006). Finally, Crb3a and b isoforms are involved in primary ciliogenesis in MDCK cells (Fan et al., 2007; Fan et al., 2004) while Crb3a is controlling the length of the hair cell kinocilia in the zebrafish otic vesicle (Omori and Malicki, 2006) suggesting that Crumbs proteins may play an important role in ciliogenesis. Besides the ciliopathy-like syndrome, a recent study has identified a *CRB2* mutation in an autosomal recessive retinitis pigmentosa (RP) (Chen et al., 2019).

In addition to *CRB2*, two other human *CRB* genes have been identified in human, *CRB1* and *CRB3* (Bazellieres et al., 2009). *CRB1* mutations are associated with several retinal degenerations from severe Leber Congenital Amaurosis (LCA) to milder RP12 (den Hollander et al., 1999) that can be mimicked by either loss of *Crb1* or *Crb2* expression in the mouse retina (Alves et al., 2013; van de Pavert et al., 2004). Lack of *Crb1* expression in the mouse retina likely results in defective junctions between Müller cells and PR and it has been suggested that these junction defects lead to PR degeneration (van de Pavert et al., 2004). There is still however the possibility that the progressive loss of PR in the retina might be a consequence of changes in signaling pathways or in apical transport since Crumbs proteins have been involved in both processes (Gurudev et al., 2014; Szymaniak et al., 2015).

All together these data suggest that CRB2 proteins might play an important role in ciliogenesis and in this study, we have uncovered a new CRB2 complex that regulate primary cilium formation that might be involved in *CRB2*-related syndrome.

## RESULTS AND DISCUSSION

### 1) CRB2 is necessary for primary ciliogenesis in ARPE19 cells

To study the role of CRB2 in primary ciliogenesis, we chose ARPE19 cells, a human retina pigmented epithelial (RPE) cell line that produces primary cilia when reaching confluency, without serum depletion (Dunn et al., 1996). We thus transiently depleted CRB2 from ARPE19 cells using siRNA transfection (Fig.1A, B, C and Fig.S1A, B) and we measured the percentage of cells with a cilium, labeled by acetylated tubulin (AcTub) antibodies, compared to control cells after 7 days (Fig.1A). Strikingly, we observed about 5 times less cilia in *CRB2* knockdown (KD) cells when compared to control cells (Fig.1B, C) clearly showing that CRB2 is required for primary ciliogenesis. Next, we tried to localize CRB2 in ARPE19 cells but all the antibodies tested gave too much non-specific background. To circumvent this difficulty we transiently expressed CRB2 tagged with the AU1 epitope and performed co-immunostainings with antibodies against AU1 and γtubulin (γTub) on cells only expressing moderate levels of AU1∷ CRB2 to avoid over-expression artifacts (Fig. 1D). We detected AU1∷CRB2 at the plasma membrane, in intracellular vesicles and, interestingly, in a cluster of vesicles nearby the centrosome, although not in cilia, that could be linked to its role in ciliogenesis (Fig. 1D).

**Figure 1:**
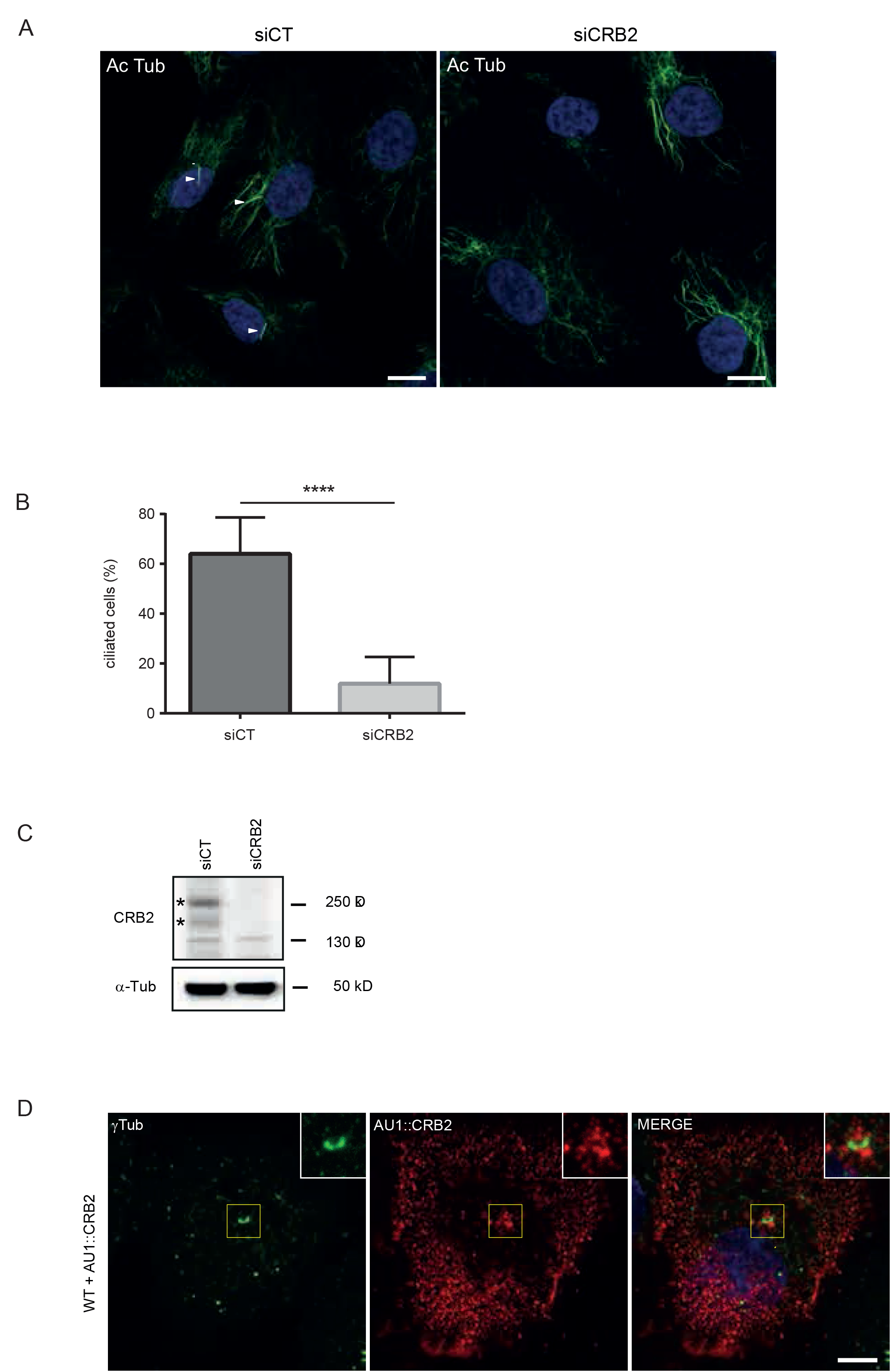
CRB2 depletion blocks ciliogenesis in ARPE19 cells. **A**: ARPE19 cells transfected with control siRNA (siCT) or a mix of two siRNA CRB2 (siCRB2) are fixed in PFA and compared for their ability to build cilia (acetylated tubulin labeling, Ac Tub) by immunofluorescence and confocal microscopy analysis. Scale bar = 10 μm. Arrowheads indicate cilia. **B**: Quantification of ciliated ARPE19 cells after transfection with siCT and siCRB2. Data are from 5 independent experiments with at least 200 counted cells per experiment per condition. **C**: Lysates from siCT or siCRB2 ARPE19 cells are analyzed by immunoblotting with antibodies against CRB2 (asterisks show specific bands) and α-tubulin (α-Tub) used as loading control (97% of CRB2 depletion in this experiment). **D:** ARPE19 cells transitorily transfected with AU1∷*CRB2* cDNA are fixed in methanol and labeled with antibodies against γ-tubulin (γTub) to localize the centrosome and against AU1 tag (to reveal exogenous CRB2). AU1∷CRB2 proteins are present in vesicles at the cellular periphery and concentrated around the centrosome which is magnified (x1.8) in the inset. Scale bar = 5 μm.

### 2) CRB2 depletion blocks the formation or the anchoring of the primary vesicle

Since ciliogenesis was impaired in *CRB2 KD* cells, we next investigated at which stage it was blocked by using ODF2 (Cenexin), a marker of the maturation of the mother centriole (MC) (Ishikawa et al., 2005) and Myosin-Va (MyoVa), a protein necessary for the attachment of the ciliary vesicles to the distal appendages of the MC (Wu et al., 2018). In *CRB2 KD* cells, ODF2 was present at levels equivalent to the ones found in control cells (Fig.2A, B) indicating that the MC has reached at least a stage of maturation before axoneme growth. TEM analysis indeed, showed that most centrioles had subdistal and distal appendages in both control and *CRB2 KD* cells (Fig.2C). We, however, noticed that most of the centrioles observed in *CRB2 KD* cells lacked a primary vesicle (Fig.2D) and the elongation of the axoneme was absent strongly suggesting that ciliogenesis was blocked at the stage of the ciliary vesicle formation or attachment. This lack of ciliary vesicle formation or attachment was not the consequence of a defect in MyoVa accumulation in the pericentriolar region (Wu et al., 2018) as we did not find a difference either in localization or in quantity of this protein between control and CRB2 KD cells (Fig.2A, B). Thus, we concluded that the defect observed in the absence of CRB2 might be the formation or the attachment of ciliary vesicle through an unknown yet mechanism to ensure their transport and anchoring to distal appendages.

**Figure 2:**
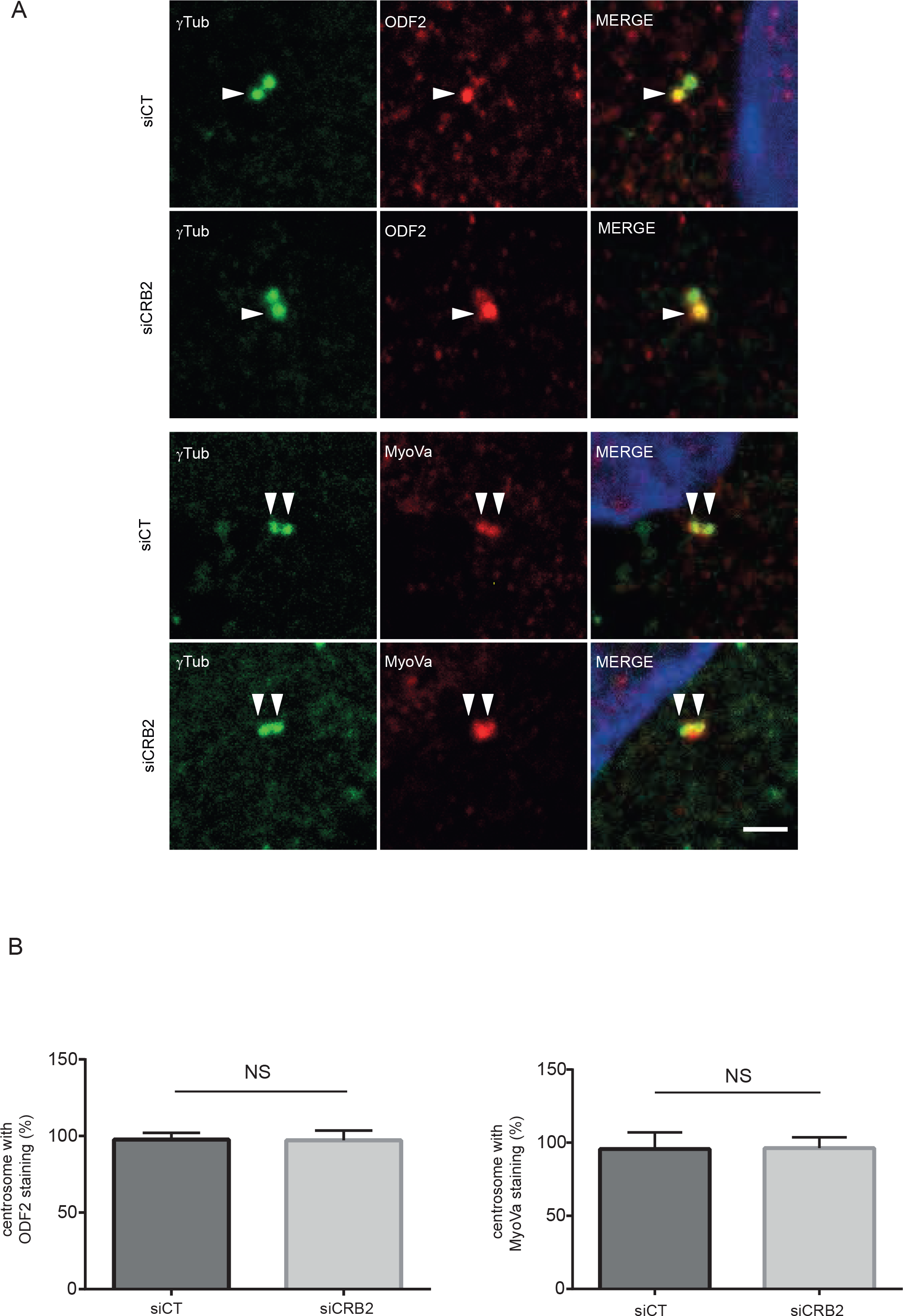
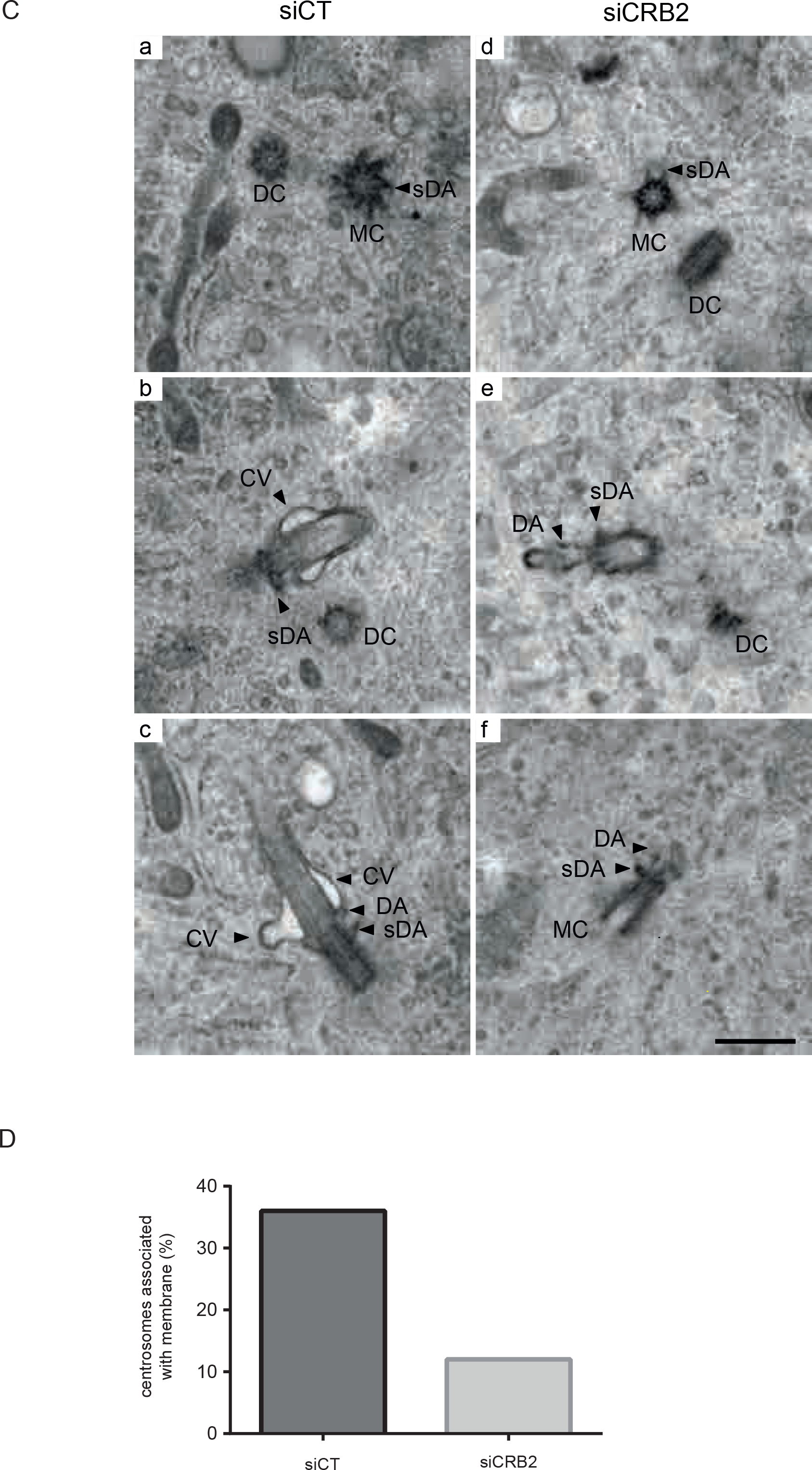
CRB2 is involved at an early stage of ciliogenesis. **A:** siCT and siCRB2 ARPE19 cells are labeled by immunofluorescence after methanol fixation with antibodies against γ-tubulin (γTub) and ODF2, or Myosin-Va (MyoVa) and visualized by confocal microscopy analysis. Scale bar = 2 μm. **B**: Percentage of centrosome labeled with ODF2 or MyoVa in siCT and siCRB2 ARPE19 cells. Results are obtained from 3 independent experiments for each protein with at least 200 (for ODF2) or 50 (for MyoVA) counted cells per experiment per condition. **C:** Both siCT and siCRB2 ARPE19 cells are assayed for primary cilium formation by TEM microscopy. a, b, c: siCT; d, e, f: siCRB2. DC: daughter centriole, MC: mother centriole, DA: distal appendages, sDA: subdistal appendages, CV: ciliary vesicle. Scale bar = 250 nm. Note the presence of CV in siCT ARPE19 cells (b,c) while it is mostly absent in siCRB2 ARPE19 cells (d,e,f). **D:** Percentage of centrosomes associated with membrane in siCT and siCRB2 ARPE19 cells. At least 100 centrosomes per condition were counted.

### 3) Ciliogenesis does not require the canonical PALS1/MUPP1 complex

Crb proteins form a complex with the cytoplasmic adapters PALS1 (MPP5) and PATJ in epithelial cells from mammals and flies (Assemat et al., 2008) while Crb1 recruits Pals1 and Mupp1 in the mouse retina (van de Pavert et al., 2004). No study, however, has yet thoroughly investigated the proteins associated with the cytoplasmic domain of CRB2. We thus used affinity-precipitation experiments with peptides mimicking the cytoplasmic domain of CRB2 to pull down potential partners in ARPE19 cells. PALS1 and MUPP1 were identified in the pull-down experiments (Fig.3A) and by mass-spectrometry of peptide-affinity pull-down (see material and methods) demonstrating that CRB2 can recruit PALS1 and MUPP1 and that these interactions relied on the presence of the C-terminal ERLI motif that binds to the PDZ domain of PALS1 (Roh et al., 2002). To test whether PALS1 or MUPP1 could play a role in primary ciliogenesis together with CRB2, we depleted these proteins in ARPE19 cells and quantified ciliogenesis as for CRB2 KD experiments (Fig. S2A, B). We found no significant effect of either protein depletion indicating that the canonical Crumbs complex does not likely play a role in impaired ciliogenesis upon CRB2 depletion. In a previous study, however, it has been shown that KD of mPals1 during early mouse cortical development led to a significant defect in ciliogenesis (Kim et al., 2010). While we found no effect of PALS1 depletion in ARPE19 cells we cannot ascertain that the amount of PALS1 remaining in depleted cells (30% in average) does not reach below a threshold necessary to block ciliogenesis (Fig.S2A). Alternatively, in the mouse study, KD of Pals1 induced a loss of mCrb2 staining suggesting that the ciliogenesis phenotype could be due to the loss of mCrb2 expression rather than a direct effect of mPals1 down-regulation. MUPP1 on the other hand was strongly depleted in our in vitro conditions (11% left in average) (Fig.S2A, B) and yet ciliogenesis was not affected further supporting that the CRB2/PALS1/MUPP1 complex might not play a crucial role during primary ciliogenesis in ARPE19 cells.

**Figure 3:**
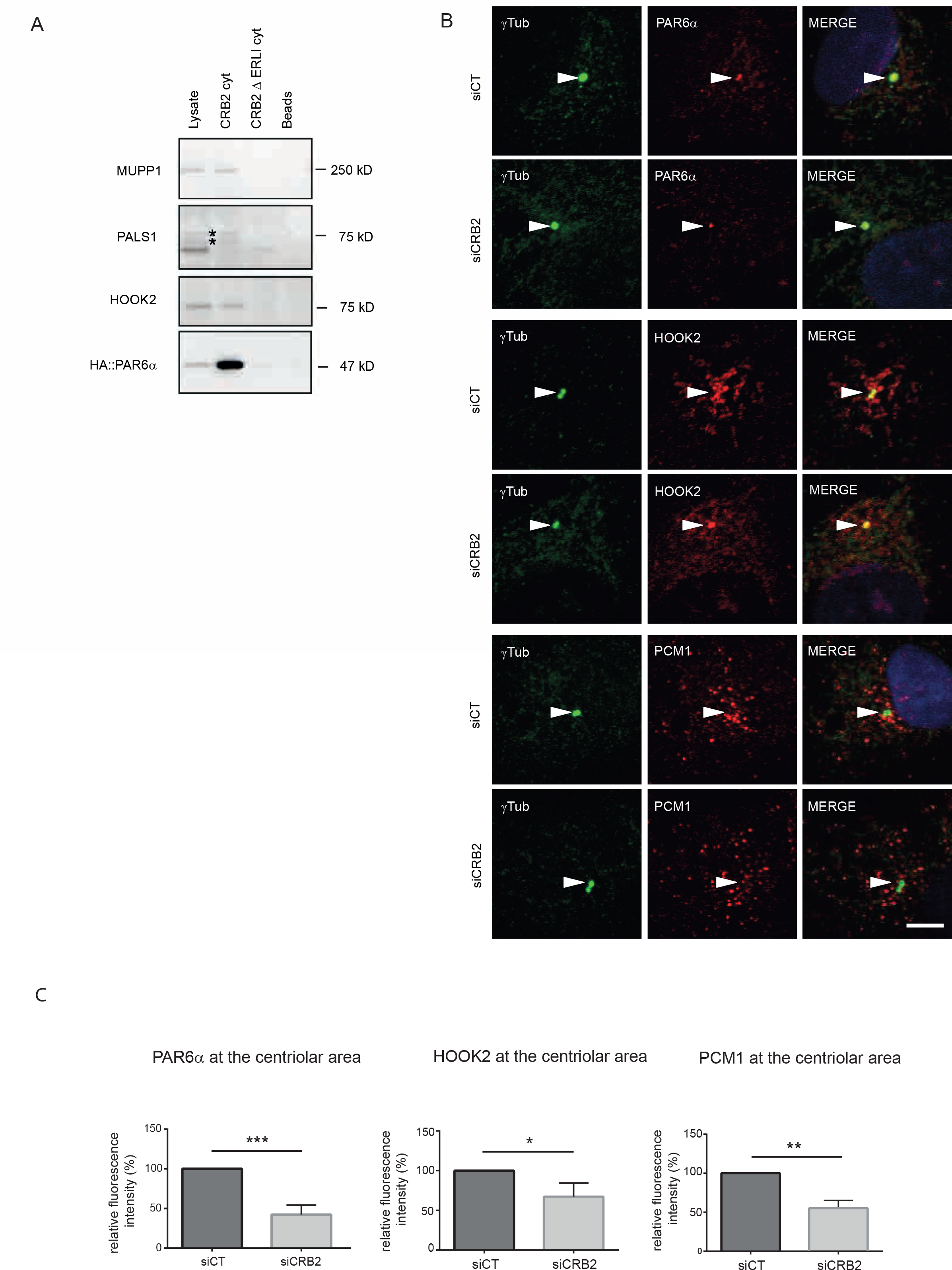
CRB2 forms a complex with PAR6α and HOOK2. **A:** HA∷PAR6α expressing ARPE19 cell lysates are used to perform peptide pull-down assays with the biotinylated cytoplasmic part of CRB2 with (CRB2 cyt) or without the last 4 amino acids ERLI (CRB2 ΔERLI cyt) bound to streptavidin beads. Lysates, peptide-pull downs and streptavidin beads alone (Beads) are processed for immunoblotting with anti-MUPP1, PALS1, HOOK2 and HA antibodies. Asterisks indicate specific bands of PALS1. **B:** siCT or siCRB2 ARPE19 cells are co-labeled after methanol fixation with antibodies against γ-tubulin (γTub) and PAR6α, or HOOK2, or PCM1, and visualized by confocal microscopy analysis. Arrowheads indicate centrosomes. Scale bar = 5 μm. **C**: Quantification of fluorescence intensity of PAR6α, HOOK2 and PCM1 at the centriolar area in siCT and siCRB2 ARPE19 cells. Data are from 3 independent experiments with at least 100 cells per experiment per condition. Values are normalized to siCT ARPE19 cells (100%).

### 4) CRB2 binds to PAR6α and HOOK2

CRB3a and Crumbs proteins bind directly to PAR6α or PAR6 both in human and drosophila respectively (Kempkens et al., 2006; Lemmers et al., 2004). In addition, we showed recently that HOOK2 directly interacts with PAR6α to recruit it to the centrosome (Pallesi-Pocachard et al., 2016) and that HOOK2, a microtubule-binding protein present in the centriolar region (Szebenyi et al., 2007a), is necessary for primary ciliogenesis in ARPE19 cells (Baron Gaillard et al., 2011). Thus, we postulated that these two proteins could be involved in CRB2 KD phenotype. To this aim, we expressed PAR6α tagged with an HA epitope in ARPE19 cells and performed CRB2 affinity pull-down experiments to test for the presence of PAR6α and HOOK2 as we did for PALS1 and MUPP1 (Fig.3A). PAR6α was specifically enriched with the CRB2 peptide containing the ERLI motif indicating that CRB2 behaves as CRB3a (Lemmers et al., 2004). We also detected HOOK2 together with PAR6α (Fig. 3A) and we thus concluded that CRB2 likely binds to HOOK2 via PAR6α. To ascertain that CRB2, PAR6α and HOOK2 form a complex in ARPE19 cells we took advantage of the fact that when mHook2 was overexpressed in ARPE19 cells, its forms membranous aggregates called aggresomes (Szebenyi et al., 2007b). When mHook2 was transiently expressed in ARPE19 cells, CRB2 and PAR6α were both recruited to mHook2 aggresomes (Fig.S3A) confirming that they can form a complex in cells. We also detected dynein which interacts with HOOK2 (Olenick et al., 2016) to transport vesicles along microtubules towards the centrosome (Szebenyi et al., 2007a). Thus, our experiments demonstrated the existence of a new CRB2 complex at least made of PAR6α, HOOK2 and dynein which is involved in the early steps of primary ciliogenesis.

### 5) CRB2 participates to the recruitment of PAR6α and HOOK2 to the centriolar region

HOOK2 and PAR6α are required for either ciliogenesis and/or centrosome organization (Baron Gaillard et al., 2011; Kodani et al., 2010) and both CRB2 (this study) and HOOK2 depletion induced a defect in the formation or anchoring of the ciliary vesicle (Baron Gaillard et al., 2011) suggesting that these proteins act together in an early stage of primary ciliogenesis. We thus decided to investigate the role of CRB2 in localizing PAR6α and HOOK2 at the centrosomal region where CRB2 positive vesicles are found close to HOOK2 and MyoVA labeling (Fig.S3B). In CRB2 KD cells, PAR6α signal was significantly decreased at the centrosome (Fig.3B, C) showing that CRB2 controls PAR6α accumulation at the centrosome. In control conditions, HOOK2 was accumulated as spots in the centriolar area and these spots were strongly diminished in the same area of CRB2 KD cells (Fig.3B,C) while PCM1 granules were more dispersed in CRB2 KD cells than in CT cells. These data demonstrate that CRB2 is necessary for the accumulation of HOOK2 and PCM1 in the centriolar area and of PAR6α at the centrosome. Together these results suggest that CRB2 present in cytoplasmic vesicles (Fig. 1D), carries PAR6α to the centrosome on microtubules towards the minus end via the dynactin subunit p150^glued^ through its interaction with PAR6α (Kodani et al., 2010) while HOOK2 enhances the interaction between dynein and dynactin to reinforce this process (Dwivedi et al., 2019a; Dwivedi et al., 2019b).

### 6) CRB2 depletion is rescued by RAB8A overexpression

In CRB2 KD as in HOOK2 KD cells, a ciliary vesicle was not formed on the activated MC (this study and (Baron Gaillard et al., 2011)). We have shown that in the case of HOOK2 depletion we could reverse the membrane transport defect by overexpressing a GFP∷RAB8A construct (Baron Gaillard et al., 2011). To test if CRB2 was acting in the same pathway, we first investigated whether CRB2 interacted with RAB8A by performing co-immunoprecipitation experiments after expression of AU1∷CRB2 and a GFP∷RAB8A in HEK cells (Fig. 4A). Indeed, AU1∷CRB2 was detected specifically albeit weakly after immunoprecipitations experiments with anti-GFP antibodies indicating that CRB2 and RAB8A belong to a common complex. We next exogenously expressed either GFP or GFP∷RAB8A in control cells and CRB2 KD cells and quantified the percentage of GFP-positive cells bearing a cilium (Fig.4B, C). Strikingly, in CRB2 KD cells transfected with GFP∷RAB8A there was a strong recovery of the ciliogenesis defect (about 5 times more than in the GFP transfected cells) clearly indicating that over-expression of GFP∷RAB8A rescued the loss of CRB2. This recovery was similar to the one observed in HOOK2 KD cells (Baron Gaillard et al., 2011), indicating that CRB2 and HOOK2 act in the same pathway together with RAB8A to build the primary cilium. Therefore, based on these data and previous already published studies, we thus propose a model for the role of this new CRB2 complex in early stages of ciliogenesis (Fig. 5). In this model CRB2 is present in vesicles that are connected to microtubules through its interaction with PAR6α and HOOK2. HOOK2 has been shown to recruit dynein and dynactin to their cargoes to increase the processivity of the complex (Dwivedi et al., 2019b) to transport CRB2 bearing vesicles towards the activated MC. In this context, CRB2 could be considered as the adaptor between the vesicles and the dynein-dynactin motor. By this mean, CRB2 also allows to concentrate HOOK2 and PAR6α to the centriolar area. In addition to this mechanism, it has been shown in *Drosophila* that Crb interacts with MyoV during the transport of Rhodopsin to the rhabdomere of PR (Pocha et al., 2011) and while we have not found a loss of MyoVa upon CRB2 depletion we cannot exclude an interaction between the two proteins during the formation of the ciliary vesicle. More work will be necessary to define the molecular mechanisms behind this step. We hope that by uncovering more of the mechanisms controlled by CRB2 during ciliogenesis will help finding more targets for improving CRB2 syndrome symptoms in patients. The fact that CRB2 and HOOK2 form a complex and that HOOK2 has been recently identified in a screen for RP in human (Patel et al., 2018) reinforces the hypothesis that these two proteins work as a complex in RP degeneration.

**Figure 4:**
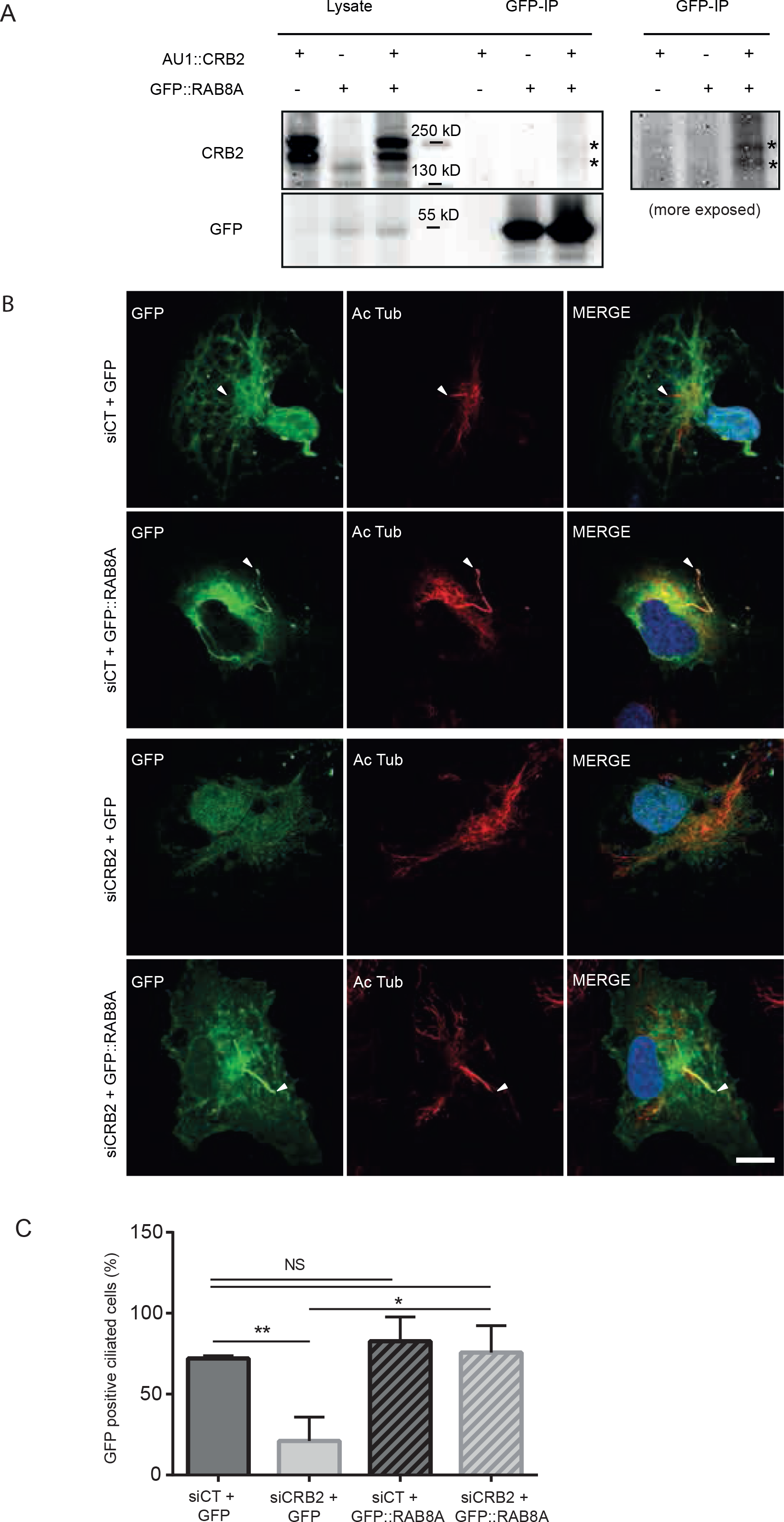
RAB8A forms a complex with CRB2 and its overexpression rescues ciliogenesis upon CRB2 depletion. **A:** Two days after transfection with AU1∷*CRB2* and/or human GFP∷*RAB8A* plasmids, HEK cells are lysed (Lysate). Cell lysates are processed for immunoprecipitation with anti-GFP antibodies (GFP-IP) which are immunoblotted with anti-CRB2 and anti-GFP antibodies. Asterisks indicate AU1∷CRB2 bands weakly co-immunoprecipitated with GFP∷RAB8A. On the right side the panel GFP-IP revealed by anti-CRB2 is more exposed to show the specificity of AU1∷CRB2 detection. **B:** siCT and siCRB2 ARPE19 cells are transfected with GFP or with GFP∷*RAB8A* cDNA. In GFP∷RAB8A siCT ARPE19 cells note the characteristic cilium overgrowth (second row) and the rescue of primary cilium in GFP∷RAB8A siCRB2 (lower row) cells. Arrowheads indicate cilia. Scale bar: 10 μm. **C:** Percentage of GFP-positive ciliated cells. siCT and siCRB2 ARPE19 cells expressing either GFP alone or GFP-*RAB8A* cDNA. Data are from 3 independent experiments with at least 70 counted cells per experiment per condition. Note that GFP∷RAB8A expression in siCRB2 cells restores ciliogenesis.

**Figure 5:**
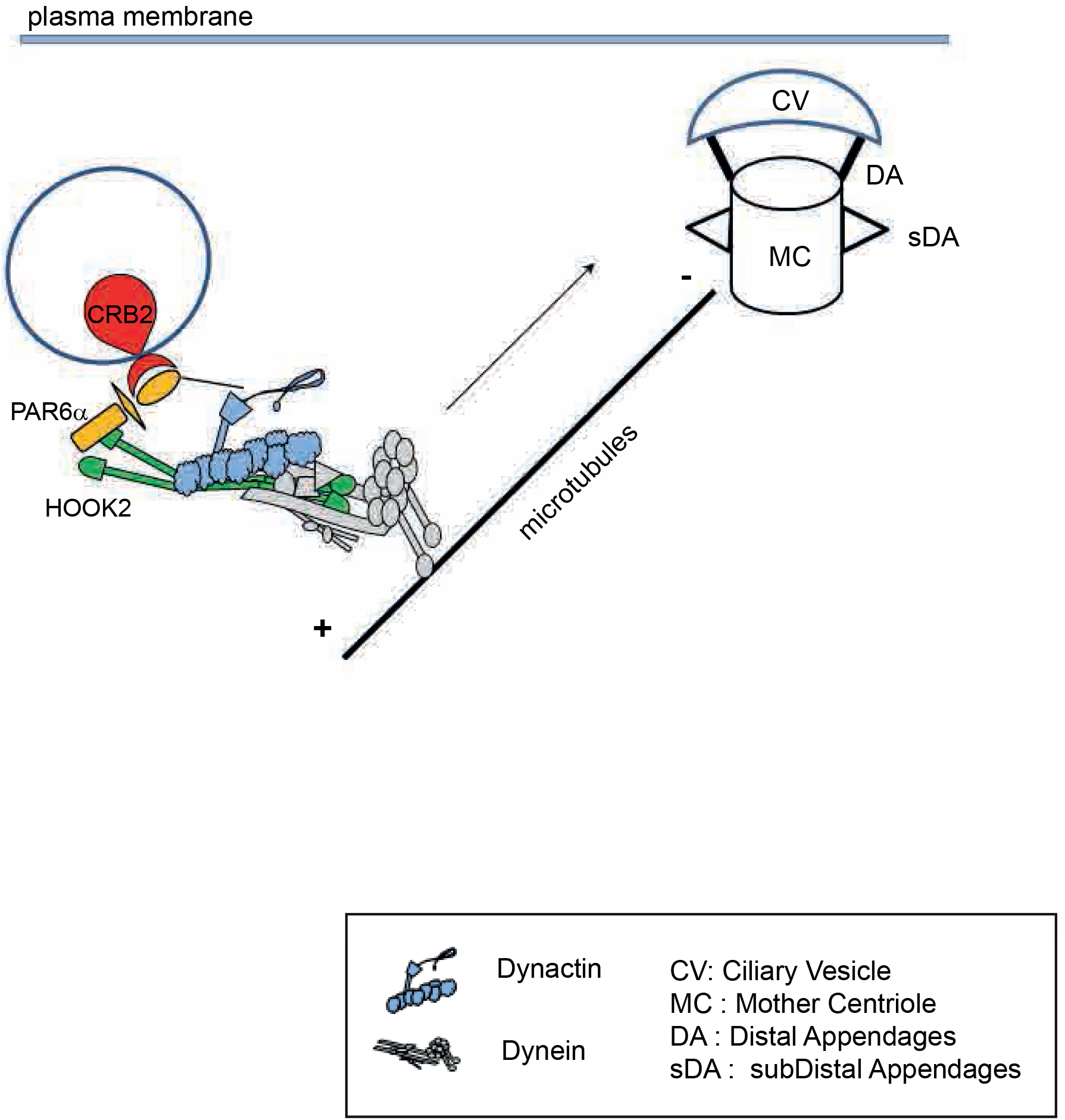
model for CRB2 role in ciliogenesis. Hypothetical model for the implication of CRB2 in the early steps of ciliogenesis. CRB2 (red) in intracellular vesicles (blue) binds to PAR6α (yellow) that in turns interacts with HOOK2 (green) which is an “activating adaptor” of dynein-dynactin complex regulating dynein mobility. In addition, PAR6α binds to dynactin subunit p150Glued and could stabilized the quaternary complex (PAR6α/Hook2/dynactin/dynein) in a motility shape. The transmembrane protein CRB2 links the PAR6α PDZ domain via its C-terminal CRB2 domain and could be an adaptor protein between vesicles and dynein-dynactin motors for transport ciliary vesicles (CV) along microtubules to the MC area.

## MATERIAL AND METHODS

### Cell culture and transfection

ARPE19 cells were cultured as in (Baron Gaillard et al., 2011). To study ciliogenesis, ARPE19 cells were cultured for 7 days in complete medium. HEK 293F cells were cultured in DMEM with L-Glutamine and 10% fetal bovine serum. ARPE19 cells were transfected with Dharmafect4 or Dharmaduo transfection reagents for siRNA or cDNA, respectively, using the manufacturer’s instructions (Horizon Discovery Dharmacon LTD, Cambridge, United Kingdom). To form aggresomes, *mHook2* cDNA was transfected in ARPE19 cells using Fugene HD (Promega France) as described in (Pallesi-Pocachard et al., 2016). HEK cells were transfected with AU1∷*CRB2* or GFP∷*RAB8A* or both together using nucleofector kit V and program A-023 of AMAXA Biosystem according to the manufacturer's instructions (Lonza Sales Ltd, Basel, Switzerland).

### siRNA

RNAi-mediated knockdown of CRB2 in ARPE19 cells was carried out using a combination of two on target plus siRNA, siCRB2 05 (J-018054-05: 5’ GCA GCU GGC CUA ACA GUAU 3’); siCRB2 07 (J-018054-07: 5’ CCU AAA CGA UGG CCA UUG G 3’) which knocked down the protein by 79% and 91% (n=3) respectively and together by 94%. Knockdown of MUPP1 (89%, n=3) and PALS1 (70%, n=3) proteins in ARPE19 cells was carried out by, respectively, siMUPP1 08 (5’ GCA CAA GUC CUU AGG CAA UUU 3’) and siPALS1 (5’ GGA UGA UGC CAA UAC GUC G 3’) siRNA were from (Horizon Discovery LTD, Cambridge, United Kingdom) except control siRNA against a sequence of luciferase (5' CGU ACG CGG AAU ACU UCG A 3') which was from Ambion (ThermoFisher Scientific, Courtaboeuf, France).

### Plasmids

*mHook2* cDNA was previously described in (Baron Gaillard et al., 2011); HA∷*PAR6α* and pEGFPC∷*RAB8A* cDNA were given by Dr I.G. Macara (University of Virginia School of Medicine, Charlottesville, VA, USA). and A. Echard (Institut Pasteur, Paris, France) respectively. AU1∷*CRB2* was designed and produced by J. Wijnholds (Department of Ophthalmology, Leiden University Medical, The Netherlands). CMV-AU1∷h*CRB2*-spA was synthesized at GenScript (USA) and cloned into pUC57, thereafter sub cloned into pUC57-ITR. pUC57-ITR-CMV-AU1∷*CRB2*-spA contained the CMV promoter, codon-optimized *CRB2* cDNA encoding a DTYRYI AU1-tag between amino acids 33/34 of *CRB2*, a synthetic poly-adenylation sequence, flanked 5’ and 3’ by inverted terminal repeats (ITR) of AAV2. pEGFP control vector was from Invitrogen (ThermoFisher Scientific, Courtaboeuf, France).

### Antibodies

The following primary antibodies were used in this study: rabbit polyclonal antibodies against-CRB2 (ThermoFisher Scientific, PA5-25628, WB: 1/200), ODF2 (Sigma-Aldrich, HPA001874, IF: 1/100), Hook2 (a gift from H. Krämer described previously in (Walenta et al., 2001), IF/WB: 1/500), MyoVa (Novus Biologicals, NBP1-92156, IF: 1/500), PAR6α (a gift from R. Lammers described previously in (Weyrich et al., 2007), IF: 1/100), PATJ (homemade, described previously in (Lemmers et al., 2002), WB: 1/300) and PCM1 (Bethyl Laboratories, A301-149A, WB/IF: 1/1000); mouse monoclonal antibodies against α-tubulin (Sigma-Aldrich, clone B-5-1-2, WB: 1/10,000), γ-tubulin (Sigma-Aldrich, clone GTU-88, IF: 1/1000), acetylated tubulin (Sigma-Aldrich, clone 6-11B-1, IF: 1/1000), cytoplasmic dynein (Covalence, MMS-400P, IF: 1/500), AU1 (Ozyme, BLE 901901, IF: 1/500), MUPP1 (BD Transduction Laboratories, clone 43, WB: 1/250), HA (Molecular Probe, A21287, WB: 1/200); chicken polyclonal antibodies against PALS1 (described previously in (van de Pavert et al., 2004) WB: 1/500) and GFP (Aves lab, GFP 1020, IF: 1/500); sheep polyclonal antibodies against GFP (University of Dundee-Scotland, WB: 1/1000) and TGN46 (AbD Serotec, AHP500G, IF: 1/1000); rat polyclonal antibodies against HOOK2 (described previously in (Baron Gaillard et al., 2011) WB: 1/200); goat polyclonal antibodies against γ-tubulin (Santa Cruz, SC7396, IF: 1/1000). Appropriate fluorochrome-conjugated or horseradish peroxidase secondary antibodies were from Jackson ImmunoResearch (Cambridge, United Kingdom) or Molecular Probes (ThermoFisher Scientific, Courtaboeuf, France), respectively.

### Immunofluorescence staining and analysis

Cultured ARPE19 cells on glass coverslips were fixed either with 3% PFA for 20 minutes and permeabilized with 1% Triton X100 for 10 minutes, or with methanol for 4 minutes at −20°C, or with ethanol 95% acetic acid 5% for 6 minutes at 4°C (for aggresome labeling). Immunofluorescence staining was processed as described previously in (Pallesi-Pocachard et al., 2016) and observed with a Zeiss (LSM 780) with X63 (1.51 NA) oil-immersion objective. Nuclei were labelled with DAPI (4’,6-diamidino-2-phenylindole). Confocal image analysis was performed using LSM image browser (Zeiss), Image J (National Institutes of Health) software. Total fluorescence of labelled PAR6α, HOOK2, PCM1 proteins was quantified inside a circle of 6 μm diameter around the centrosome (γTub labelled) using a macro command under Image J software.

### Transmission electron microscopy

siRNA treated ARPE19 cells were cultured on glass coverslips and treated as in (Baron Gaillard et al., 2011). Serial sections containing centrosomes were selected and observations were performed with an electron microscope FEI Morgagni (FEI, Netherlands) operating at 80 kV. Micrographs were taken with an Olympus SIS Mega view III digital camera (Olympus, Japan).

### Western Blot

For Western blot analysis, cells were lyzed in 50 mM Tris-HCL pH 7.4, 150 mM NaCl, 1% Triton X-100 (buffer A) with protease inhibitor cocktail containing 1 μg/ml antipain, 1 μg/ml pepstatin, 15 μg/ml benzamidine, and 1 μg/ml leupeptin. The cell lysates were centrifuged at 100,000g during 30 min and the supernatants were completed with Laemmli buffer and βmercaptoethanol. The proteins were heated and processed by SDS-PAGE, transferred to nitrocellulose, stained with Ponceau red, and immunoblotted with indicated antibodies. The proteins were detected with the Western lightning chemi-luminescence reagent plus (Perkin Elmer) and visualized by MyECL imager (ThermoFisher Scientific, Courtaboeuf, France).

### Co-immunoprecipitations

HEK cells were transfected with AU1∷*CRB2* or GFP∷*RAB8A*, or GFP∷*RAB8A* and AU1∷*CRB2* cDNA. After 48h, cells were treated as in (Pallesi-Pocachard et al., 2016). Each pre-cleared lysate was incubated with GFP-Trap A beads (Chromotek, Germany) overnight at 4°C. After 4 washes with buffer A, bound proteins were analyzed by Western blotting after SDS-PAGE.

### Peptide pull-down assays

ARPE19 cells transiently transfected to express HA∷*PAR6α* were treated as in the previous section. 150 μL pelleted streptavidin beads (streptavidin agarose resin, ThermoFisher Scientific, Courtaboeuf, France) were coated with 2mg of a biotinylated peptide mimicking the cytoplasmic part of CRB2 (CRB2 cyt: amino-acids 1249 to 1285) or without the four terminal amino-acids which constitute a PDZ-binding domain (CRB2 ΔERLI cyt: amino-acids 1249 to 1281) both synthetized by CovaLab (Cambridge, UK). 15 μl of streptavidin beads with either biotinylated peptides were incubated with HA∷PAR6α ARPE19 lysate at 4°C overnight. After four washes with buffer A, bound proteins were analyzed by Western blotting after SDS-PAGE. For mass spectrometric experiments, peptides corresponding to carboxy-termini of CRB2 (amino-acids 1271 to 1285) and CRB2 ΔERLI (amino-acids 1271 to 1281) were synthesized (CovaLab, Cambridge, UK) with an additional lysine at the N-terminus and cross-linked to NHS-activated Sepharose 4 Fast Flow beads (GE Healthcare). Pull-down experiments were performed using ARPE cell lysates and interacting proteins were identified by mass spectrometry after in-gel digestion as described in (Mojallal et al., 2014).

### Statistics

Statistical analysis were performed with GraphPad Prism version 6.00 and based on an unpaired T-test analysis. ****: P<0.0001; ***: 0.0001<P<0.001; **: 0.001<P<0.01; *: 0.01<P<0.05; NS (no significant difference): P>0,05.

## ACKNOWLEDGEMENTS

We thank the Le Bivic Lab members for discussions during lab meetings and critical reading of the manuscript and Helmut Kramer, University of Texas, USA, for the gift of rabbit anti-HOOK2 antibodies. We acknowledge the PiCSL-France BioImaging core facility of IBDM supported by the Agence Nationale de la Recherche (ANR-10-INSB-04-01, call "Grand Emprunt"). This project was supported by CNRS and Aix-Marseille University (UMR7288) and the labex INFORM (grant ANR-11-LABX-0054). The authors declare no competing financial interests.

## AUTHOR CONTRIBUTION

MBR and DMH: design and realization of the experiments. MBR: validation, visualization and statistical analysis. SA and JPB: Mass Spectrometry experiments and analyses. MBR, DMH, JW and ALB: conceptualization and writing of the manuscript.

## Abbreviation list

AJs: Adherens Junctions
aPKC: atypical Protein Kinase C
ARPE: Adult Retinal Pigmented Epithelia
CRB: CRumBs
HEK: Human Embryonic Kidney
LCA: Leber Congenital Amaurosis
MDCK: Madin Darby Canine Kidney
MPP5: Membrane Palmitoylated Protein 5
MUPP1: MUlti-PDZ domain Protein-1
MyoVa: Myosin-Va
ODF2: Outer Dense Fiber protein 2
PALS1: Protein Associated with Lin Seven
PAR6: PARtitioning-defective 6
PATJ: PALS-1-Associated Tight Junction
PCM1: PeriCentriolar Material 1
CV: Ciliary Vesicle
PR: PhotoReceptor
RP: Retinis Pigmentosa
RPE: Retinal Pigmented Epithelia
TEM: Transmission Electron Microscopy

---

**Supplementary Figure 1:**
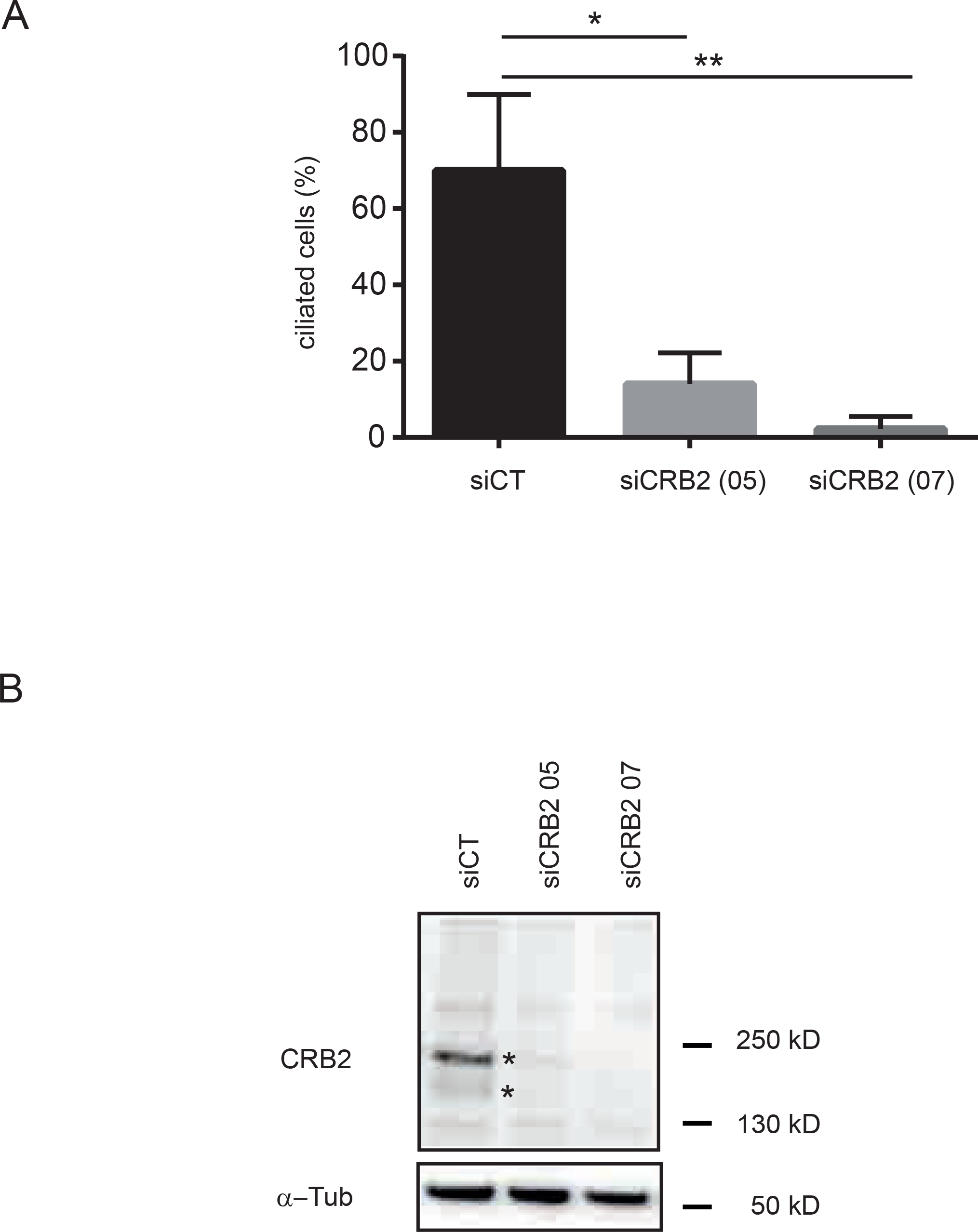
Effect of CRB2 siRNAs. **A**: Quantification of ciliated cells in siCT, siCRB2 05 and siCRB2 07 ARPE19 cells. Data are from 3 independent experiments with at least 150 counted cells per experiment per condition. **B:** Lysates from siCT, siCRB2 05 and siCRB2 07 ARPE19 cells are analyzed by immunoblotting with antibodies against CRB2 (asterisks show specific bands) and α-tubulin (α-Tub) used as loading control. 89% (siCRB2 05) and 93% (siCRB2 07) of CRB2 depletion in this experiment.

**Supplementary Figure 2:**
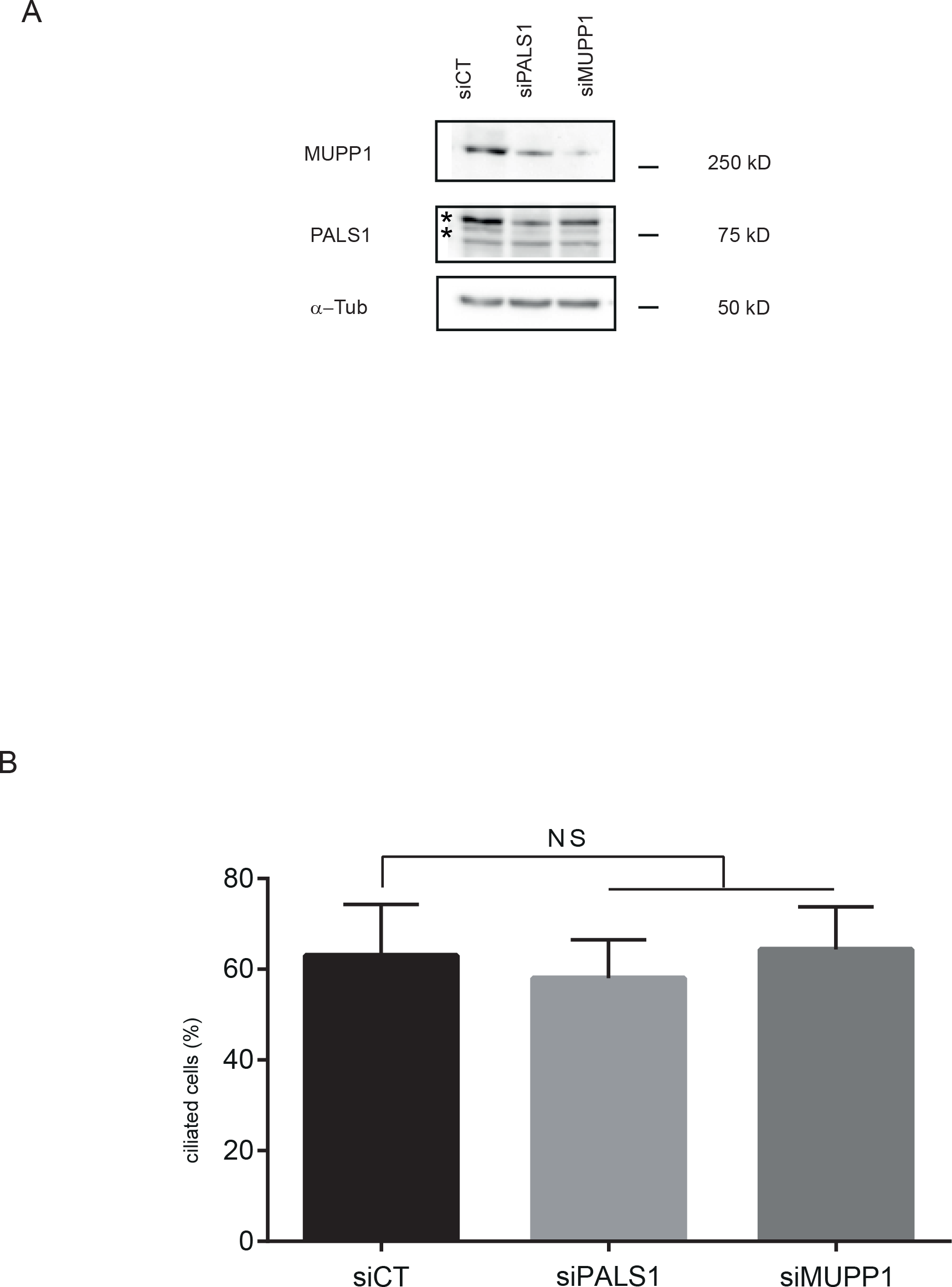
The canonical CRB complex is not involved in ciliogenesis in ARPE19 cells. **A:** Lysates from siCT, siPALS1 and siMUPP1 ARPE19 cells are analyzed by immunoblotting with antibodies against MUPP1, PALS1, and α-tubulin (α-Tub) used as loading control. 67% (siPALS1) and 89% (siMUPP1), of respectively PALS1 or MUPP1 depletion in this experiment. **B:** Percentage of ciliated cells in siCT, siPALS1 and siMUPP1 ARPE19 cells. Data are from 3 independent experiments with at least 140 counted cells per experiment per condition.

**Supplementary Figure 3:**
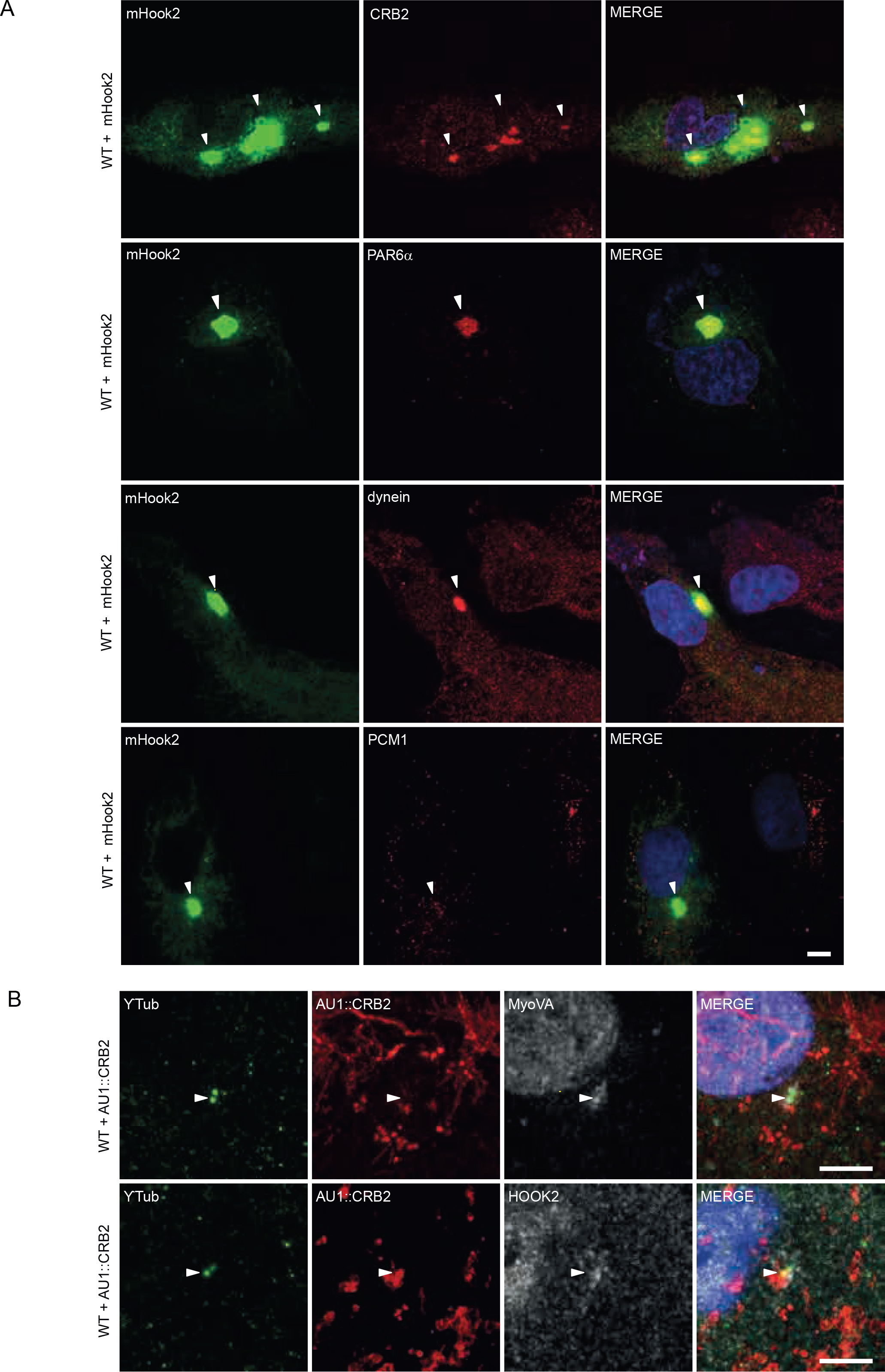
CRB2 and HOOK2 colocalization. **A**: ARPE19 cells are fixed by ethanol/acetic acid after transient overexpression of mouse Hook2 (mHook2) for forty hours. Overexpression of mHook2 promotes the accumulation of CRB2, PAR6α and dynein in aggresome but not PCM1. Arrowheads indicate aggresomes. Note that endogenous aggregated CRB2 was detected in aggresomes. Scale bar = 10 μm. **B:** ARPE19 cells transiently transfected with AU1∷*CRB2* cDNA are fixed in methanol and labeled with antibodies against γ-tubulin (γTub) and AU1 tag, and antibodies against MyoVA or HOOK2. Arrowheads indicate centrosome. Scale bar = 5 μm.

